# Derivation of ringed seal (*Phoca hispida*) induced multipotent stem cells

**DOI:** 10.1101/2020.05.20.105890

**Authors:** Violetta R. Beklemisheva, Polina S. Belokopytova, Veniamin S. Fishman, Aleksei G. Menzorov

**Affiliations:** Institute of Molecular and Cellular Biology of the Siberian Branch of the Russian Academy of Sciences, Novosibirsk, Russia; Institute of Cytology and Genetics of the Siberian Branch of the Russian Academy of Sciences, Novosibirsk, Russia; Novosibirsk State University, Novosibirsk, Russia

**Keywords:** reprogramming, pluripotency, Carnivora, iPS cells, Rex1, adipocyte differentiation

## Abstract

Induced pluripotent stem (iPS) cells have been produced just for a few species among order Carnivora: snow leopard, Bengal tiger, serval, jaguar, cat, dog, ferret, and American mink. We applied the iPS cell derivation protocol to the ringed seal (*Phoca hispida*) fibroblasts. The resulting cell line had the expression of pluripotency marker gene *Rex1*. Differentiation in embryoid body-like structures allowed us to register expression of *AFP*, endoderm marker, and *Cdx2*, trophectoderm marker, but not neuronal (ectoderm) markers. The cells readily differentiated into adipocytes and osteocytes, mesoderm cell types of origin. Transcriptome analysis allowed us to conclude that the cell line does not resemble human pluripotent cells, and, therefore, most probably are not pluripotent. Thus, we produced ringed seal multipotent stem cell line capable of differentiation into adipo- and osteocytes.

## INTRODUCTION

Order Carnivora consists of two suborders: Caniformia and Feliformia. It includes domestic animals, that are used for disease modeling, fur-bearing animals, such as mink, marine mammals, and other species. Derivation of induced pluripotent stem (iPS) cells could allow insights into pluripotency and embryonic development of these species, as well as the development of new disease models. Currently, pluripotent cells were produced from several Carnivora species: dog (Luo et al., 2011; Baird et al., 2015; Lee et al., 2011; Vaags et al., 2009; Shimada et al., 2010; Whitworth et al., 2012; Koh et al., 2012; Tsukamoto et al., 2018), snow leopard (Verma et al., 2012), Bengal tiger, serval, jaguar, and cat (Verma et al., 2013; Dutton et al., 2019; Gómez et al., 2010), ferret (Gao et al., 2020), and American mink (Menzorov et al., 2015).

Marine mammals represent the basal Carnivora group. As their embryos are unavailable, iPS cells would expand our knowledge of embryonic development and differentiation. We attempted to generate ringed seal (*Phoca hispida*) iPS cells. We were able to produce multipotent stem cells that expressed pluripotency and mesenchymal stem cell marker *Rex1* and were able to differentiate them into mesoderm cell types, adipo- and osteocytes.

## MATERIALS AND METHODS

### Production of ringed seal fibroblasts

Primary fibroblasts of the ringed seal were obtained from lung necropsy. Tissue samples from wild female were collected during aboriginal quota sealing in the coastal waters of the Bering Sea (Mechigmen bay, Chukotka Autonomous Okrug, Russia). To establish primary fibroblast cell culture, we used a conventional technique (Stanyon, Galleni, 1991). The fibroblast culture medium consisted of a-MEM supplemented with 15 % fetal bovine serum (FBS), 1x MEM non-essential amino acids solution, 1x GlutaMAX supplement, and 1x penicillin-streptomycin (Thermo Fisher Scientific, USA).

### Production of ringed seal multipotent stem cell line

To reprogram the ringed seal fibroblasts we used lentiviral vectors LeGO (http://www.lentigo-vectors.de/vectors.htm) with *EGFP* and human reprogramming transcription factors: *OCT4, SOX2, C-MYC* and *KLF4*, courtesy of Dr. Sergei L. Kiselev, Moscow. We used the previously published protocol (Menzorov et al., 2015) with minor modifications. Lentiviruses were produced in the Phoenix cell line using Lipofectamine 3000 (Thermo Fisher Scientific, USA). The multiplicity of infection was estimated as 5.1 using *EGFP* lentiviral vector (Beklemisheva, Menzorov, 2018). Fibroblasts at passage 4 (3 × 10^5^ cells, 30 × 10^3^ cells/cm^2^) plated the day before were transduced with viruses containing four reprogramming transcription factors: 50 % virus supernatant, 50 % fibroblast culture medium without antibiotics with heat-inactivated FBS, and 10 µg/ml Polybrene. Transduction was performed for two consecutive days. Cells were passaged onto a 6 cm cell culture dish coated with 0.1 % gelatin on mouse strain CD-1 feeder cells on day 5. From day 6 we used iPS cell culture medium: a-MEM supplemented with 20 % ES cell qualified FBS, 1x MEM non-essential amino acids solution, 1x GlutaMAX supplement, 0.1 mM 2-mercaptoethanol, 1x penicillin-streptomycin, and 10 ng/ml bFGF recombinant human protein (Thermo Fisher Scientific, USA). From day 6 until day 12 the medium was changed once in two days with the addition of 1 mM valproic acid (Sigma-Aldrich, USA). On day 23 colonies were picked up and expanded on the feeder. The passage was performed with 0.25 % Trypsin-EDTA (Thermo Fisher Scientific, USA). All cell cultures were maintained at 37°C and 5 % CO_2_.

Multipotent stem cell derivation was performed at the Collective Center of ICG SB RAS “Collection of Pluripotent Human and Mammalian Cell Cultures for Biological and Biomedical Research” (http://ckp.icgen.ru/cells/; http://www.biores.cytogen.ru/icg_sb_ras_cell/).

All animal studies were undertaken with prior approval from the Ethics Committee on Animal and Human Research of the Institute of Molecular and Cellular Biology SB RAS, Russia (protocol No. 01/20 of 11 February 2020).

### Cytogenetic analysis

Cytogenetic analysis for fibroblasts was carried out on passage 7 and for multipotent stem cell line on passage 9. The preparation of metaphase chromosomes from fibroblasts was performed as previously described (Yang et al., 1999; Graphodatsky et al., 2000; Graphodatsky et al., 2001). GTG-banding of metaphase chromosomes was done according to a previously published protocol (Seabright, 1971). For each cell line, averages of 50 conventionally stained by Giemsa metaphase plates were analyzed. Digital images were captured using the VideoTest system (Zenit, St. Petersburg, Russia) with a charge-coupled device (CCD) camera (Jenoptik, Jena, Germany) mounted on a Zeiss microscope Axioscope 2 (Zeiss, Oberkochen, Germany). Metaphase spreads images were edited in Corel Paint Shop Pro Photo X2 (Corel, Ottawa, Canada). Chromosomes of the ringed seal (*P. hispida*) were arranged according to the current nomenclature (Graphodatsky et al., 2020).

### Multipotent stem cell differentiation

Differentiation into embryoid body-like (EB-like) structures was performed in EB differentiation medium, the same composition as iPS cell culture medium but with 20 % FBS instead of ES qualified FBS and without bFGF. Cells were passaged into 1 % agarose coated cell culture plates to prevent attachment. The medium was changed every second day for 31 days. On day 5 part of EBs were plated onto 0.1 % gelatin-coated cell culture plates for osteocyte and adipocyte differentiation. Osteocyte differentiation was carried out in EB differentiation medium; adipocyte differentiation in a similar adipocyte differentiation medium supplemented with 10 % knockout serum replacement (KSR) (Thermo Fisher Scientific, USA) instead of FBS from day 6.

### Cytochemical staining

Cells were fixed by 4 % paraformaldehyde for 20 min and washed with PBS. Adipocytes were stained with 0.7 % Sudan black B in propylene glycol, washed twice with 0.85 % propylene glycol, and washed multiple times with PBS. Calcification was shown by staining with alizarin red (Sigma-Aldrich) according to the manufacturer’s recommendations. Staining was analyzed on Zeiss Observer.Z1 fluorescent microscope with AxioCam HRm 3 CCD-camera (Zeiss, Germany). Digital images were analyzed using the ZEN 2 starter (Zeiss, Germany) software.

### DNA isolation

Genomic DNA was isolated from cells using a PCR buffer with nonionic detergents (PBND), which was adapted from a protocol from Perkin Elmer Cetus (Higuchi, 1989).

### RNA isolation and cDNA synthesis

RNA was isolated using Aurum Total RNA mini kit (Bio-Rad, USA). Genomic DNA was removed using *DNase*I (Fermentas, USA), 0.4 micrograms of total RNA were used for cDNA synthesis by First Strand cDNA Synthesis kit (Thermo Fisher Scientific, USA).

### Primer design and PCR

We used Primer-BLAST software (Ye et al., 2012) to design primers for mink *Rex1* and canine *Cdx2*: qNvRex1F 5’-AAA GCG TTT TCC ACA CCC CT-3’, qNvRex1R 5’-CTC CTT GTC CAT GGT CCT CG-3’, CfCdx2E1F 5’-GGA ACC TGT GCG AGT GGA TG-3’, and CfCdx2E3R 5’-TTC CTT TCC TTG GCT CTG CG-3’.

We used previously published primer sequences for human-specific *KLF4* transgene (Mathew et al., 2010), *Mycoplasma* detection (Choppa et al., 1998), mink *Hprt1* (Rouvinen-Watt et al., 2012), mink *Oct4* (Menzorov et al., 2015), and human *AFP* (Huangfu et al., 2008).

PCR was performed using BioMaster HS-Taq PCR-Color (2×) (Biolabmix, Russia) in 10 µL reaction volume.

### *De novo* transcriptome assembly and dataset annotation

We performed non-stranded & polyA-selected mRNA library preparation and PE100 sequencing on DNBSEQ at BGI (People’s Republic of China). The number of reads was 62,390,350, Q20 rate – 95.77 %, and GC rate – 50.78 %. The raw sequence data are available in NCBI BioProject repository, accession number: PRJNA718133, link: https://www.ncbi.nlm.nih.gov/bioproject/PRJNA718133.

RNA-seq read quality was assessed using FastQC (https://www.bioinformatics.babraham.ac.uk/projects/fastqc/). The RNA was isolated from cell culture containing mouse feeder cells. To exclude reads originating from feeder transcripts we aligned reads to the mouse genome using bowtie2 (Langmead, Salzberg, 2012) with default parameters. All aligned reads were excluded from the following analysis. *De novo* transcriptome assembly was done with Trinity (v. 2.11.0) using default settings (Grabherr et al., 2011; Haas et al., 2013). Coding regions of the assembled transcripts were predicted using TransDecoder (v. 5.5.0) with default settings (Haas et al., 2013). We used the Basic Local Alignment Search Tool (BLAST) and the Swiss-Prot database to assign functional annotations (Altschul et al., 1990; Camacho et al., 2009). We assessed the quality of transcriptome assembly using BUSCO (Seppey et al., 2019) and Trinity scripts. The following filtration was done using homemade python scripts. We used StringTie for generating transcript quantifications with default options (Pertea et al., 2015). RNA-seq data analysis was performed using https://usegalaxy.org/ server (Afgan et al., 2018) and Computational Cluster of the Novosibirsk State University (Russia).

### Analysis of endo- and exogenous expression of *OCT4, KLF4, SOX2*, and *c-MYC* genes

We extracted all ringed seal transcripts encoding *OCT4, KLF4, SOX2*, and *c-MYC* using BLAST against the Swiss-Prot database. For this analysis, we used transcriptome assembly obtained from unfiltered reads (i.e. possibly containing mouse feeder transcripts), to allow capturing of transcripts conserved between mouse, human, and ringed seal. We found that each of the target genes was represented by multiple Trinity transcripts. We manually annotated all obtained transcripts using BLAST nucleotide alignment and BLAST protein alignment against NCBI nucleotide or protein collections. For three genes (except *Sox2*), we found that some transcripts showed better alignment to Carnivora orthologues, whereas other transcripts had better alignment to primates orthologues. These results were consistent between nucleotide and protein alignments and therefore allowed us to classify transcripts by organism of origin.

To qualitatively estimate expression levels of the target genes we used bowtie2 (Langmead, Salzberg, 2012) to align reads to assembled transcripts of ringed seal and human orthologues. All reads aligned to both human and ringed seal references were removed, and the remaining read counts were used as a proxy of gene expression.

### Genome-wide comparison of gene expression between cell types and species

We obtained gene expression quantification files from ENCODE using batch download with a filter (organism: human; assay: RNA-seq; file format: tsv). All files were processed using homemade Python scripts to obtain FPKM values. We filtered out genes that do not have one-to-one orthologues between mouse, human, and ringed seal. For the remaining genes, we only kept the top 10% of genes that showed the highest FPKM standard deviation values across samples. The resulting gene expression matrix (1843 genes) was used to compute Spearman’s R correlation coefficient between each pair of samples.

## RESULTS

### Multipotent stem cell derivation and differentiation

The experiment outline is shown in Figure 1A. We aimed to produce iPS cells from primary ringed seal fibroblasts and used the following human reprogramming transcription factors: OCT4, SOX2, C-MYC, and KLF4. On day 26 after transduction, there were colonies with different morphology. Two colonies were picked up and expanded (Fig. 1B). One of them, iPHIS1, gave rise to cells that grew in a monolayer (Fig. 1C,D), the other had fibroblast morphology after the passage and was discarded. Overgrown culture formed “bubbles” (Fig. 1E) that later detached and floated as cyst-like structures resembling canine embryos (Hayes et al., 2008). The cells continue to divide after passage 15.

**Fig. 1.**
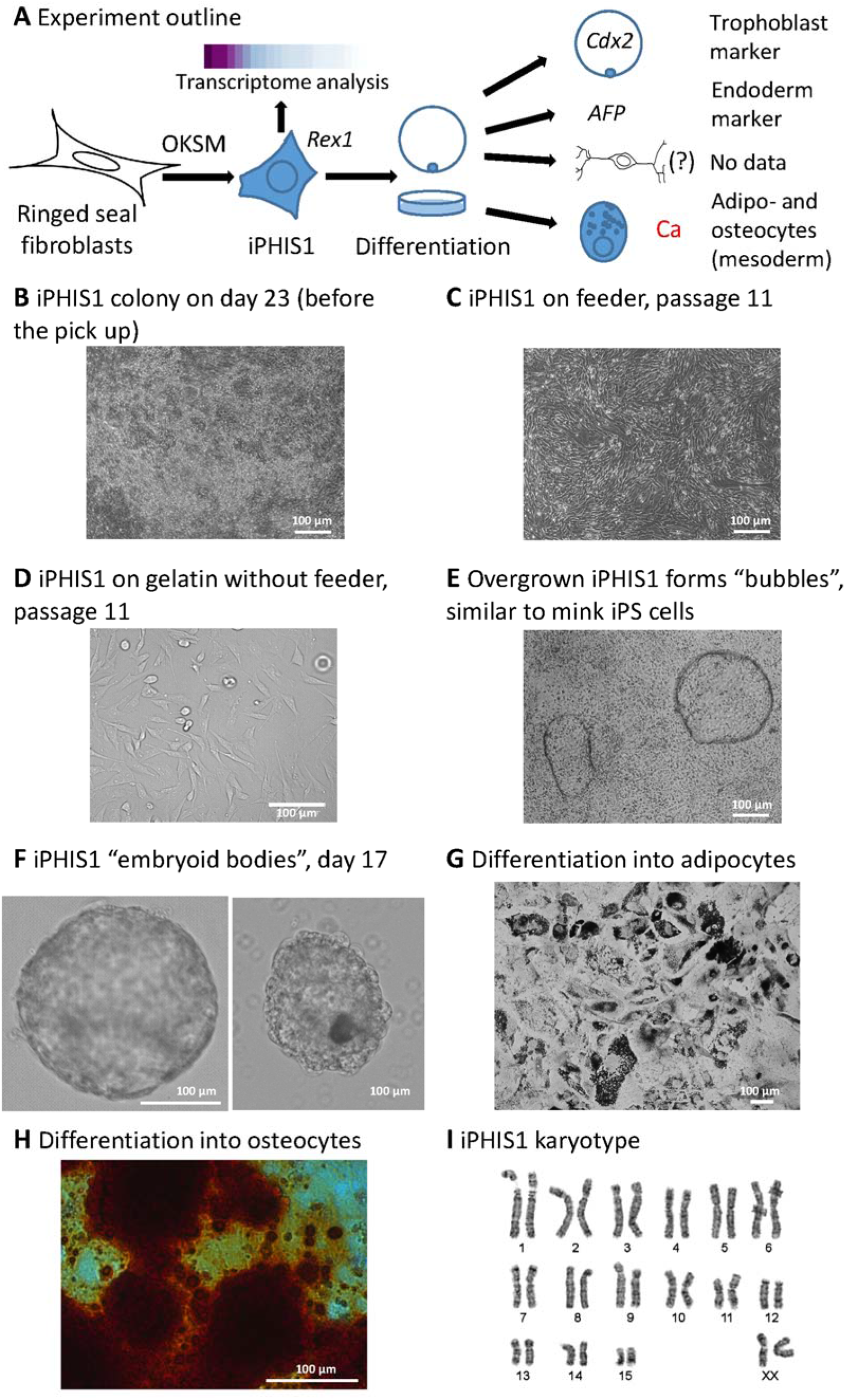
Derivation, colony morphology, differentiation, and karyotype of ringed seal induced multipotent stem cells.

We attempted to differentiate iPHIS1 cells into derivatives of the three germ layers. Solid and cyst-like EB-like structures were formed after passage to non-adhesive culture plates (Fig. 1F). Their gene expression analysis is described below. Adipo- and osteogenic differentiation was successfully performed in the adipo- and EB differentiation media, respectively (Fig. 1G,H). Interestingly, there was some calcification in the adipocyte differentiation medium (data not shown).

### Cytogenetic analysis

We performed the cytogenetic analysis of the ringed seal (*P. hispida*) primary fibroblasts and iPHIS1 cells. Fibroblasts had 32 chromosomes, typical for this species (Arnason, 1974), XX (n = 50) (Fig. 1I) with 15.7 % polyploid cells. The iPHIS1 cells (n = 50) had 32 (n = 47), 33 (n = 2), and 35 (n = 1) chromosomes, with 5 % polyploid cells. We had not revealed a difference in the pattern of GTG-banding between karyotypes of the primary fibroblasts and iPHIS1 cells. We conclude that the iPHIS1 karyotype is stable. A propensity for the emergence of tetraploid cells in pinniped fibroblast cultures has been noted previously (Árnason. 1974). We also observed a certain proportion of tetraploid cells in primary fibroblast cell lines established from different pinniped species (Beklemisheva et al., 2020. This feature depended on the passage number, but the species karyotypes were found to be stable in diploid cells.

### Gene expression pattern

First, we applied RT-PCR analysis of gene expression. We were able to show the expression of one of the key pluripotency markers *Rex1* (*Zfp42*) as well as *Oct4* in iPHIS1 cells by RT-PCR (Fig. 2A). Analysis of EB-like structures revealed the expression of *AFP*, endoderm marker, and *Cdx2*, trophoblast marker (Fig. 2B). We also analyzed DNA samples of ringed seal fibroblasts and iPHIS1 cells; they were negative for *Mycoplasma* contamination.

**Fig. 2.**
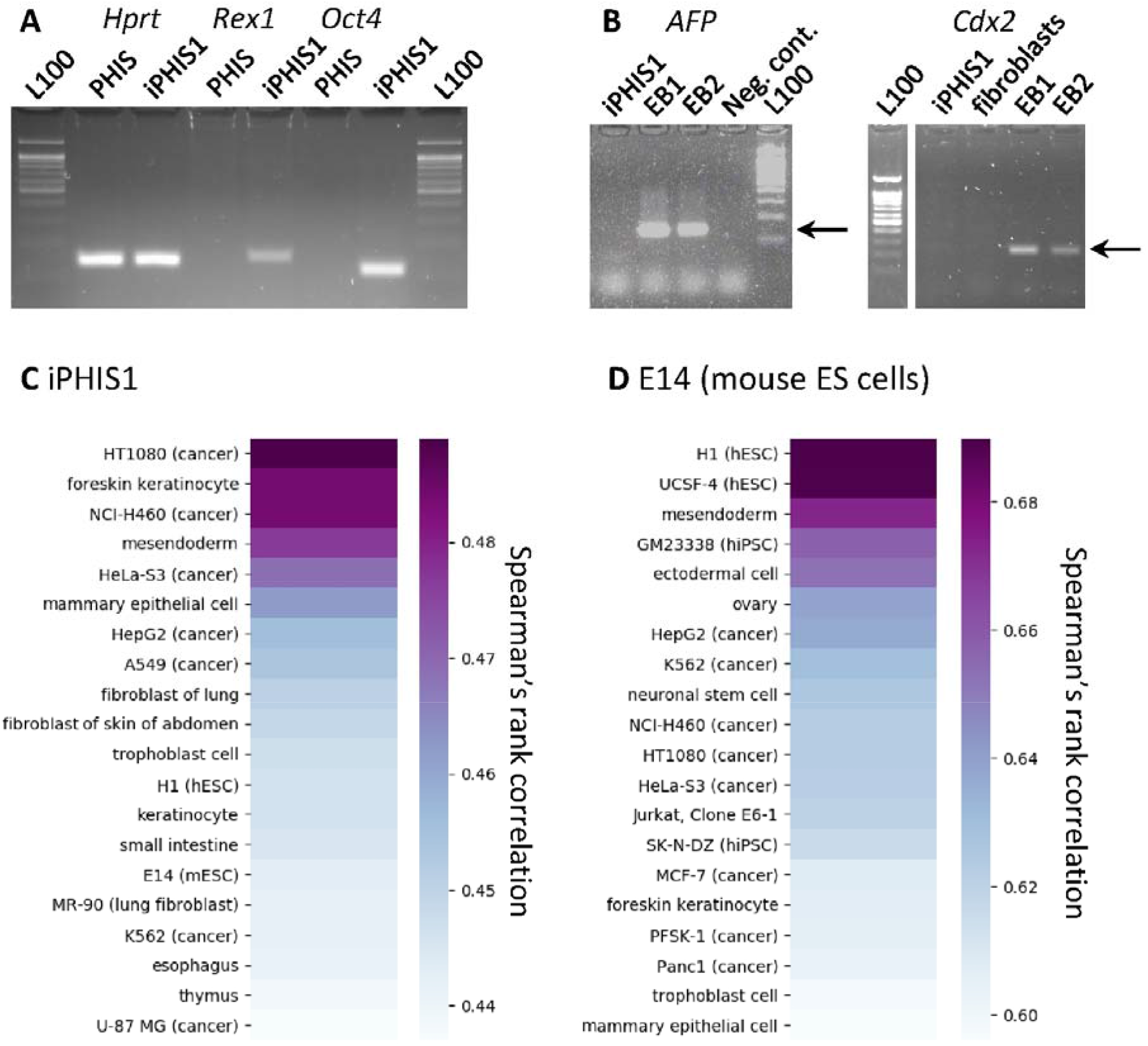
Gene expression in iPHIS1 and Spearman’s rank correlation coefficients between iPHIS1 expression pattern and human samples. (A) Gene expression in iPHIS1 and PHIS fibroblasts. (B) Gene expression in iPHIS1 embryoid body-like structures. (C) Top 20 Spearman’s correlation coefficients of iPHIS1 and human ENCODE data with addition of mouse E14 ES cell line. (D) Top 20 Spearman’s correlation of mouse ES cell line E14 and human ENCODE data.

We next employed transcriptome sequencing to profile genome-wide gene expression in iPHIS1 cells. To avoid contamination originating from mouse feeder cell transcripts, we filtered out all reads aligned to the mouse genome (8.5 % of reads) and subjected the remaining dataset to *de novo* transcriptome assembly. This resulted in a draft assembly composed of 170,758 genes (209,671 transcripts) (http://dx.doi.org/10.17632/5mkvk5yc4w.1). BUSCO orthologues analysis showed that 67 % out of the Mammalia BUSCO database were present in this draft assembly, confirming its high quality.

We then used the Swiss-Prot protein database to annotate obtained Baikal seal genes, which allowed us to find orthologues for 31,194, containing 12,228 unique gene names (http://dx.doi.org/10.17632/5mkvk5yc4w.1).

We used obtained transcriptomic data to profile the expression of some marker genes in iPHIS1 cells (http://dx.doi.org/10.17632/5mkvk5yc4w.1). We confirmed a relatively high level of *Rex1* expression. There was not *Nanog* in the Trinity transcripts. We aligned reads to *P. vitulina Nanog* and were able to find just 44 reads. We presume that those transcripts represent a slight DNA contamination, but not the *Nanog* expression. As we introduced human *OCT4, KLF4, SOX2*, and *c-MYC* genes to reprogram fibroblasts into iPS cells, we decided to check whether those genes are expressed or silenced. Lentiviral transgenes are silenced in pluripotent stem cells, thus their silencing would suggest passing through pluripotency. We performed an analysis of endo- and exogenous expression of their transcripts and found expression of both human transgenes and ringed seal orthologues of *Oct4* (*Pou5f1*), *Klf4*, and *c-Myc*. As for *Sox2*, we found that Trinity failed to assemble a ringed seal orthologue. Manual analysis of reads aligned to *P. vitulina Sox2* reference showed the presence of ringed seal transcripts, although expressed at the relatively low level compared to exogenous human *SOX2* gene. Consistent with that, all isoforms of the *Sox2* gene assembled by Trinity showed 100% identity to the human *Sox2* gene. Thus, there were no signs of transgene silencing.

Next, we performed a genome-wide comparison of expression patterns observed in iPHIS1 cells and various human cell types with added mouse ES cell line E14. For this analysis, we used all available human RNA-seq data from ENCODE, 288 samples. We accessed the similarity of gene expression profiles using Spearman’s correlation coefficient (Fig. 2C). To test the applicability of this approach, we performed the same analysis comparing mouse ES cell line E14 transcriptome data with the same human cell lines and tissues (Fig. 2D). As expected, mouse ES cells showed the highest similarity to human pluripotent cells. In contrast, iPHIS1 expression shows the highest correlation with several cancer cell lines, as well as with foreskin keratinocytes, but not with pluripotent cells. These data argue against the establishment of pluripotency in iPHIS1 cells.

## DISCUSSION

Transcription factors Oct4, Sox2, c-Myc, and Klf4 were used to generate iPS cells from a variety of Carnivora species. We decided to produce iPS cells from the ringed seal (*P. hispida*), the basal representative of the suborder Caniformia. Only one cell line expressed pluripotency marker genes *Rex1* and *Oct4*. Its morphology differed from mouse, human, and mink iPS cells (Fig. 1C,D). Similar to American mink iPS cells, overgrown cell culture formed “bubbles” (Fig. 1E) that later unfastened and floated as cysts. We differentiated iPHIS1 cells into EB-like structures to analyze differentiation potential (Fig. 1F). Cyst-like EBs resembled canine embryos (Hayes et al., 2008), expressed endoderm marker *AFP* and trophoblast marker *Cdx2* (Fig. 2B). EBs plated on gelatin efficiently differentiated into mesoderm derivatives: adipo- and osteocytes. We were not able to show ectoderm differentiation, as *Tubb3* marker gene was expressed in both iPHIS1 and EBs (data not shown). The presence of *Tubb3* transcript in undifferentiated pluripotent stem cells is in accordance with our previous data on mouse ES cells (Menzorov et al., 2019). Ability to differentiate into adipo- and osteocytes allows us to presume that iPHIS1 cells are at least multipotent.

The expression of a trophoblast marker upon differentiation, *Cdx2*, was rather unexpected, though trophoblast differentiation was shown for human and mouse primed pluripotent stem cells (Xu et al., 2002; Kojima et al., 2014) and canine iPS cells (Luo et al., 2011; Wilcox et al., 2008). If iPHIS1 cells were pluripotent, differentiation into trophoblast would suggest a primed pluripotency state.

We used bFGF to derive and propagate the iPHIS1 cell line. Colonies with prospective morphology were not formed in the media supplemented with LIF or combination of LIF and bFGF. Canine pluripotent stem cells were produced with LIF (Luo et al., 2011; Baird et al., 2015; Lee et al., 2011; Vaags et al., 2009) or LIF and bFGF (Shimada et al., 2010; Whitworth et al., 2012; Koh et al., 2012; Tsukamoto et al., 2018). Also, cells were later cultured in the presence of LIF only (Whitworth et al., 2012). Other Carnivora species pluripotent cells include snow leopard (Verma et al., 2012), Bengal tiger, serval, jaguar, and cat (Verma et al., 2013; Dutton et al., 2019) cultured with LIF; cat cultured with LIF and bFGF (Gómez et al., 2010), and ferret cultured with bFGF (Gao et al., 2020). Interestingly, cat iPS cells required species-specific feline bFGF (Dutton et al., 2019), thus species-specific LIF or bFGF may be beneficial. We generated American mink iPS cells without LIF of bFGF supplementation (Menzorov et al., 2015), though inactivated mouse embryonic fibroblasts secret both growth factors. Different requirements of the iPS cell culture of various species indicate that different signaling pathways are activated. It leads to different pluripotency states, naïve, primed, or other, as additional distinct pluripotency states had been described recently. More high-quality transcriptome data may facilitate the distinction between various pluripotency states in different species.

Pluripotent stem cells of different pluripotency status also have distinctive morphology. For instance, American mink pluripotent stem cells form colonies unlike mouse and human naïve and primed cells (Menzorov et al., 2015). Thus, morphology and expression of several marker genes are not enough to determine whether cells are pluripotent or not.

Transcriptome analysis of iPHIS1 cells revealed that their expression pattern resembled some human cancer cell lines, keratinocytes, and mesodermal cells (Fig. 2C). Correlation of expression pattern of iPHIS1 and human pluripotent cells H1 and mouse E14 was slightly lower, although within the top quartile of all correlation coefficient values observed. Thus, transcriptome analysis allows concluding, that iPHIS1 cells do not resemble human embryonic stem cells, as pluripotent stem cells were not among the closest by correlation. At the same time, this analysis was able to place mouse pluripotent cells close to human ones, thus this approach gives meaningful results.

Our data allows presuming that iPHIS1 cells are not pluripotent. The question remains, whether this cell line was produced by fibroblast reprogramming. If there was a pluripotent state, transgene silencing would be expected. Retroviral transgene silencing in pluripotent stem cells is a well-known phenomenon (Maherali et al., 2007; Wernig et al., 2007; Okita et al., 2007). The transcriptome data revealed expression of *Oct4, Klf4*, and *c-Myc* human transgenes comparable with their endogenous counterparts. As for Sox2, the endogenous expression was lower than that of the transgene. There are three main explanations. First, iPHIS1 cells were reprogrammed in an intermediate pluripotency state and/or to iPS cells and subsequently differentiated into multipotent stem cells. The pluripotency state stage was short and not enough to silence the lentiviral transgenes. Second, we directly reprogrammed ringed seal fibroblasts to multipotency. It was recently shown, that *Oct4, Klf4, Sox2*, and *c-Myc* can reprogram somatic cells to a variety of cell types, including stromal ones (Schiebinger et al., 2019). Third, there is a possibility that a mesenchymal stem cell in a fibroblast population gave rise to iPHIS1, probably even without reprogramming. It would explain the adipo- and osteogenic differentiation of iPHIS1, as well as *Oct4* and *Rex1* expression. As for *AFP* and *Cdx2* gene expression (endoderm and trophoblast markers, respectively), their expression may be a culture artefact or a property of mesenchymal stem cells in a given culture conditions, i.e. during EB-like differentiation. Also, there are some other cell types that can differentiate into adipo- and octeocytes, such as pericytes.

We produced only one ringed seal multipotent stem cell line, experiments for iPS ?ell derivation were not successful. Suboptimal culture conditions might be the main reason. ES cell qualified FBS batch are only tested for mouse ES cell growth support. Also, growth factor origin may be important. We used human bFGF, and for cat iPS cells only species-specific feline LIF was able to support pluripotency (Dutton et al., 2019). Growth factor concentration is an important factor as well. Ferret iPS cells were derived with human bFGF, but in 10x concentration compared to human iPS cells and this study (Gao et al., 2020). Different researchers used small molecules to enhance iPS cell derivation efficiency. In our experience, 2i inhibitors (PD0325901 and CHIR99021) and TGF-β antagonist A83-01 caused substantial fibroblast death, thus we were not able to use them. Another way to increase efficiency is to use different transgenes and/or delivery vectors. We have recently shown that Sendai virus -based vector transgene expression level is superior to lentiviruses (Beklemisheva, Menzorov, 2018). Also, another set of reprogramming factors was successfully used to reprogram not only human, but ferret fibroblasts to pluripotency, OCT4, SOX2, KLF4, L-MYC, LIN28A, mp53DD, and 160 oriP/EBNA-1 (Gao et al., 2020).

## CONCLUSIONS

We produced ringed seal multipotent stem cells, capable of differentiation into mesoderm cell types, adipo- and osteocytes. We were able to show expression of endoderm (*AFP*) and trophoblast (*Cdx2*) markers after differentiation, but not ectodermal. Transcriptome analysis suggests that iPHIS1 cells do not resemble human or mouse pluripotent stem cells. Their differentiation profile suggests mesenchymal stem cell identity, though we do not have sufficient data to prove it by transcriptome analysis. The iPHIS1 could be used as a model cell line for ringed seal adipo- and osteogenesis studies.

## ACKNOWLEDGMENT

The reported study was funded by RFBR, project number 20-04-00369, and the Ministry of Education and Science of the Russian Federation, state project 0259-2021-0016.

Authors are thankful to Michael Zelensky, Alexey Ottoj, and the Community of the Chukotka Autonomous Region indigenous “Lorino” (Russian Federation) for assistance in the ringed seal tissue sample collection. The authors gratefully acknowledge the primary fibroblast cell culture of *P. hispida* provided by the “Molecular and Cellular Biology” core facility of the Institute of Molecular and Cellular Biology SB RAS (project 0310-2018-0011).

## AUTHOR CONTRIBUTIONS

VRB obtained ringed seal fibroblasts and performed cytogenetic analysis. AGM produced and differentiated multipotent stem cells, performed RT-PCR gene expression analysis. PSB performed transcriptome analysis. VSF supervised transcriptome analysis. AGM carried out interpretations of the data and project coordination; AGM did most of the writing with contributions from VRB, PSB, and VSF. All authors read and approved the final manuscript.

